# Genome-wide multi-layered epigenomic profiling across human aging

**DOI:** 10.64898/2026.07.27.740934

**Authors:** Mara Steiger, René Krüger, Mina Shaigan, Deepika Puri, Giulia Fornero, Hannes Klump, Alexander Meissner, Ivan G. Costa, Helene Kretzmer, Wolfgang Wagner

## Abstract

Aging is characterized by highly reproducible alterations across multiple layers of the epigenetic landscape, including DNA methylation, chromatin accessibility, and histone modifications. However, it remains unclear to what extent these age-associated changes are interconnected and coordinated coherently. To investigate the genome-wide distribution and interplay of age-associated epigenetic alterations, we generated whole-genome bisulfite sequencing (WGBS) data from blood samples of 120 healthy donors. Integration with ATAC-seq data revealed no clear relationship between age-related changes in DNA methylation and chromatin accessibility. We further examined the association of these alterations with age-dependent changes in CTCF occupancy and histone modifications, including H3K27ac, H3K27me3, H3K4me1, H3K4me3, and H3K9me3, but observed very little corresponding changes in chromatin states. Collectively, our integrative genome-wide analysis revealed only limited association between age-associated epigenetic alterations in DNA methylation, chromatin accessibility, and histone modifications, arguing against a broadly coordinated remodeling of the aging epigenome.

## Introduction

Epigenetic alterations are a hallmark of aging and provide robust biomarkers for estimating chronological age. The first epigenetic clocks were developed from DNA methylation (DNAm) profiles ^1,2^, and similar predictors have since been established based on chromatin accessibility ^3,4^ and histone modifications ^5^. Recent efforts have largely focused on improving the accuracy of such epigenetic predictors and enhancing their ability to capture biological rather than chronological aging ^6^. Considerably less is known about how these distinct layers of the aging epigenome are related to one another. Their association has often been postulated, since histone modifications are major determinants of chromatin accessibility, while several histone marks recruit DNA methyltransferases (DNMTs) to specific genomic regions, thereby shaping DNAm patterns ^7–9^. Whether these established molecular interactions translate into coordinated age-associated changes across the epigenome, however, remains unresolved. Clarifying this relationship could provide important insights into why age-associated epigenetic alterations preferentially arise at specific genomic loci and their underlying mechanisms.

A comprehensive integration of age-associated DNAm with other epigenetic layers requires genome-wide datasets. The vast majority of human methylome studies rely on Illumina BeadChip arrays, which provide highly reproducible and cost-effective profiling across large cohorts ^10^. However, even the latest MethylationEPIC v2.0 array interrogates fewer than 4% of the approximately 28 million CpG sites in the human genome, leaving most intergenic regions and repetitive elements unexplored. Whole-genome bisulfite sequencing (WGBS) overcomes these limitations but has been applied only rarely in human aging studies. Existing WGBS datasets are typically constrained by limited sequencing depth coverage ^11^, small cohort sizes ^12^, or heterogeneous tissues ^13^. Consequently, a systematic genome-wide characterization of age-associated DNAm in blood – the tissue most widely used for epigenetic clock studies – has remained lacking.

## Results

### Whole genome analysis of age-associated DNA methylation

We performed WGBS of peripheral blood samples from 120 healthy donors aged 19–78 years (Supplementary Fig. 1a; Supplemental Table 1). Principal component analysis (PCA) of genome-wide DNA methylation (DNAm) profiles revealed a gradual age-dependent separation of samples (Fig. 1a). Global mean DNAm showed a significant negative correlation with chronological age (R = -0.50; Fig. 1b), consistent with the long-standing concept of age-associated global hypomethylation ^14^. However, our genome-wide analysis demonstrates that this decline is remarkably modest, corresponding to a decrease of only 0.024 percentage points in mean DNAm per year. Even CpGs flanked by an A or T (‘W’) on both sides (WCGW tetranucleotides), which are considered particularly susceptible to stochastic methylation loss during aging ^15^, exhibited only a moderate reduction in methylation (Supplementary Fig. 1b).

**Figure 1.**
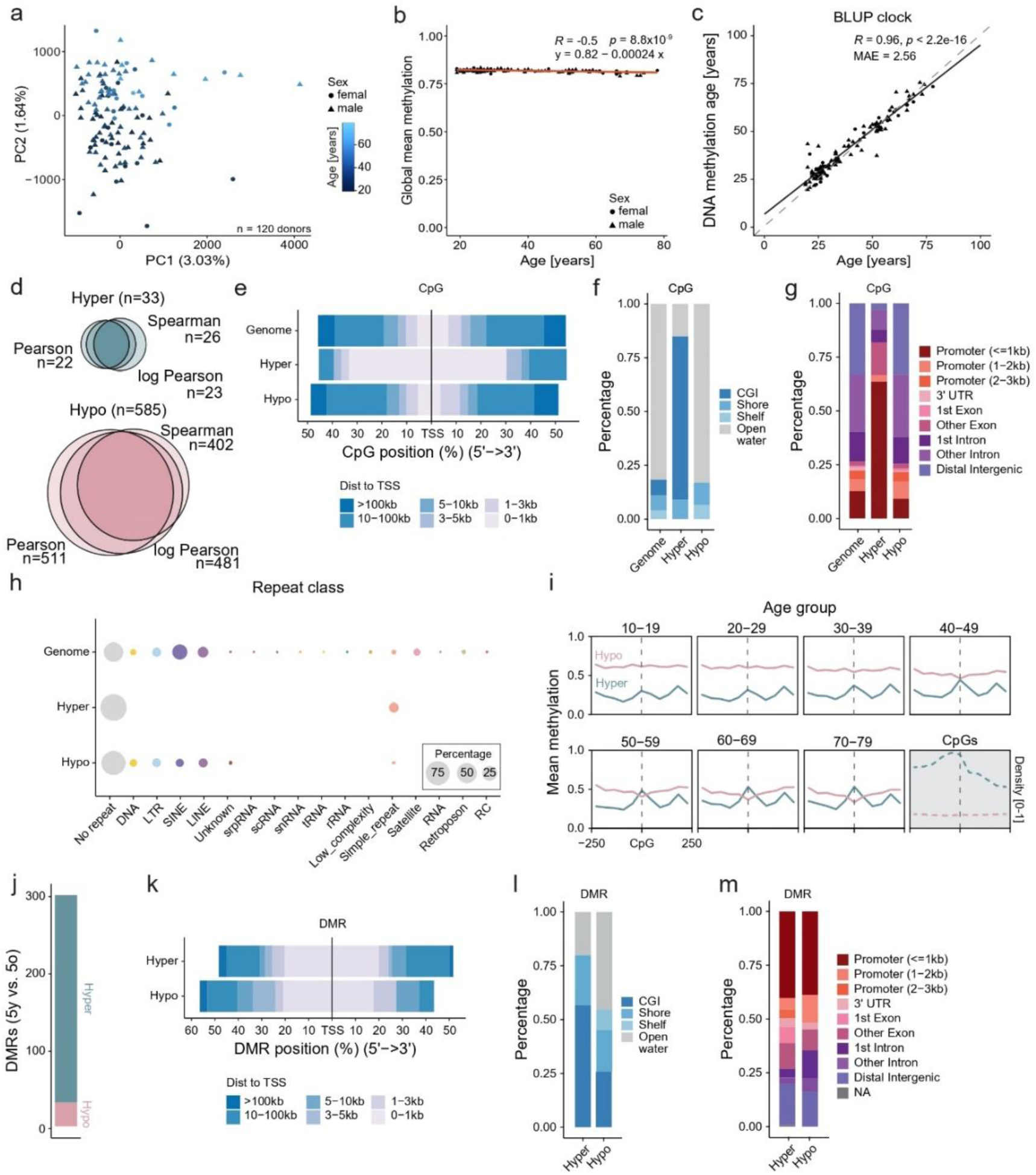
Genome-wide analysis of age-associated DNAm in human blood. **a)** Principal component analysis (PCA) of WGBS profiles from 120 donors reveals a gradual separation of samples according to chronological age. **b)** Mean genome-wide DNAm levels across all CpGs decrease modestly with age (β = –0.00024; *P* = 8.8 × 10^-9^). **c)** Epigenetic age predictions generated using the BLUP clock ^26^. Pearson’s correlation coefficient (*R*) and the median absolute error (MAE) between predicted and chronological age are indicated. **d)** Age-associated CpGs were identified using Pearson, log-transformed age Pearson, or Spearman correlation with chronological age (|*R*| > 0.6; adjusted *P* < 0.05). The three analytical approaches showed substantial overlap in the identified CpGs. **e–g)** Age-hypermethylated CpGs were significantly enriched **e)** at transcription start sites (TSSs), **f)** within CpG islands (CGIs), and **g)** in promoter regions. **h)** Strongly age-hypermethylated CpGs were preferentially located in non-repetitive genomic regions. **i)** Age-hypomethylated CpGs exhibited broad methylation loss across neighboring CpG sites, whereas age-hypermethylated CpGs showed localized gains in DNA methylation, forming peaks spanning approximately 150 bp. **j)** Differentially methylated regions (DMRs) were identified by comparing five young and five old donors (≥10 CpGs within 300 bp; adjusted *P* < 0.05). **k–m)** Age-associated DMRs were enriched at **k)** transcription start sites. DMRs gaining DNA methylation with age showed modest enrichment in **l)** CpG islands and **m)** promoter regions.

Because repetitive elements comprise a substantial fraction of genomic CpGs, they contribute disproportionately to global methylation levels. Previous studies have suggested pronounced age-associated hypomethylation of repetitive elements, particularly short interspersed nuclear elements (SINEs), including Alu elements, and to a lesser extent long interspersed nuclear elements (LINEs) and long terminal repeat (LTR) retrotransposons ^16–18^. In contrast, our WGBS data revealed only modest methylation losses that were comparable to the genome-wide average across (Supplementary Fig. 1c). Together, these findings support the concept of global age-associated hypomethylation but indicate that the overall loss of methylation is surprisingly small and occurs to a similar extent in repetitive and non-repetitive genomic regions.

We next assessed whether established array-based epigenetic clocks could accurately estimate age from our WGBS data. Although such clocks often perform suboptimally on bisulfite sequencing datasets because of incomplete coverage of clock CpGs ^19^, all clocks tested – including the Horvath clock ^20^, Skin & Blood clock ^21^, Hannum clock ^22^, Han clock ^23^, PhenoAge ^24^, Weighted kernel density estimation (WKDE) ^25^, the elastic net based clock (EN clock) ^26^, and the best linear unbiased prediction (BLUP clock) ^26^ – showed significant correlations between predicted and chronological age (Supplementary Fig. 1d). Prediction accuracy generally increased with the number of CpGs included in the model. The BLUP clock, which incorporates 319,607 CpGs after quality control, achieved the highest performance (Fig. 1c; R = 0.96; mean absolute error = 2.56 years). These results demonstrate that our high-coverage WGBS dataset robustly captures age-associated methylation patterns and is well suited for comprehensive analyses of epigenetic aging.

### Focus on age-correlated CpGs and genomic regions

Age-associated DNAm changes do not necessarily follow linear trajectories across the lifespan and may instead exhibit logarithmic or other monotonic patterns ^27^. To capture different potential age-dependent relationships, we therefore identified age-associated CpGs using three complementary approaches: Pearson correlation, log-transformed age Pearson correlation, and Spearman rank correlation (always with absolute correlation > 0.6, false discovery rate (FDR)-adjusted *p* value < 0.05, and coverage > 10 in at least 50% of samples), which identified 533, 504, and 428 significant CpGs, respectively (Supplemental Table 2). The three approaches showed substantial overlap and their combined union comprised 618 CpGs, including 585 age-hypo and 33 age-hypermethylated sites (Fig. 1d), which were distributed across the genome (Supplementary Fig. 2a,b). Among the age-associated hypermethylated sites were established epigenetic aging markers, including CpGs within *ELOVL2* and *FHL2* (Supplementary Fig. 2c), which have previously been reported as robust age-associated loci ^23,28^. The correlation strengths observed in our WGBS dataset were lower than those previously reported using Illumina BeadChip arrays or targeted methylation assays, likely reflecting differences in measurement platforms with lower coverage in some WGBS samples.

We next investigated the genomic distribution of age-associated CpGs. The relatively few age-hypermethylated CpGs were enriched at transcription start sites (TSSs), within CpG islands (CGIs), and in promoter regions, whereas hypomethylated CpGs displayed a genomic distribution largely comparable to the background distribution of all CpGs (Fig. 1e–g). Analysis of repeat element annotations further demonstrated that hypermethylated CpGs were predominantly located in non-repetitive genomic regions, while hypomethylated CpGs showed a repeat element distribution similar to the genomic background (Fig. 1h).

Age-associated hypomethylated CpGs were accompanied by a broad reduction in DNAm across the surrounding region. In contrast, hypermethylated CpGs displayed a more localized methylation increase centered around the identified CpG, with flanking enrichment patterns extending approximately 150 bp followed by neighboring peaks (Fig. 1i). This pattern may reflect regional regulatory organization, potentially related to nucleosome positioning or coordinated regulation within CpG-rich genomic contexts.

We subsequently identified age-associated differentially methylated regions (DMRs), which may better capture coordinated regional methylation changes than individual age-associated CpGs. Comparing five young (21–26 years) and 5 older donors (66–78 years), which were subsequently also used for chromatin state analysis, we identified 268 hypermethylated and 31 hypomethylated DMRs (Fig. 1j; Supplementary Fig. 2d; Supplemental Table 3). Unlike individual age-associated CpGs, age-associated DMRs showed a relatively higher proportion of hypermethylated regions and a distinct genomic distribution. Hypomethylated DMRs were preferentially enriched at TSSs, whereas the enrichment of hypermethylated DMRs within CGIs and promoter regions was less pronounced compared with individual hypermethylated CpGs (Fig. 1k–m).

We next examined whether age-associated DNAm alterations were reflected in transcriptional changes using a public RNA-sequencing dataset ^3^. Only a small subset of individual age-associated CpGs was located near genes exhibiting significant age-related expression changes (FDR-adjusted *p* < 0.01), and overall, no consistent relationship between methylation directionality and gene expression changes was observed (Supplementary Fig. 2e). Similarly, none of the identified age-associated DMRs showed significant associations with age-related differential gene expression (Supplementary Fig. 2f). The limited correspondence between age-associated DNAm alterations and transcriptional changes is consistent with previous reports and may reflect the relatively small magnitude of methylation changes occurring during aging, as well as the complex relationship between DNA methylation and transcriptional regulation ^29,30^.

### Age-associated changes in chromatin accessibility

To investigate whether age-associated DNAm changes are linked to alterations in chromatin accessibility, we generated ATAC-seq profiles from peripheral blood mononuclear cells (PBMCs) of 96 individuals from the same cohort. After quality control, 91 samples were retained for downstream analyses (Supplementary Fig. 3a,b). Peak calling identified 239,599 open chromatin regions (OCRs). We next identified age-associated changes in chromatin accessibility using the same correlation-based framework applied to DNAm analysis: OCRs were selected based on Pearson correlation, log-transformed age Pearson correlation, or Spearman rank correlation with chronological age. Because accessibility changes showed smaller effect sizes compared with DNAm alterations, we applied a less stringent correlation threshold of |R| ≥ 0.25 while maintaining an FDR-adjusted *p* value < 0.05 (Supplementary Fig. 3c). The three approaches identified largely overlapping sets of age-associated OCRs, resulting in a combined set of 12,424 opening and 21,300 closing regions (Fig. 2a,b, Supplemental Table 4). Despite their statistical association with age, the magnitude of accessibility changes was generally modest (Fig. 2c). Age-associated closing OCRs were preferentially located near TSS and promoter regions, whereas opening OCRs were predominantly enriched in distal intergenic regions (Fig. 2d,e), consistent with previous reports describing age-associated remodeling of promoter and enhancer-associated chromatin accessibility ^3,4^.

**Figure 2.**
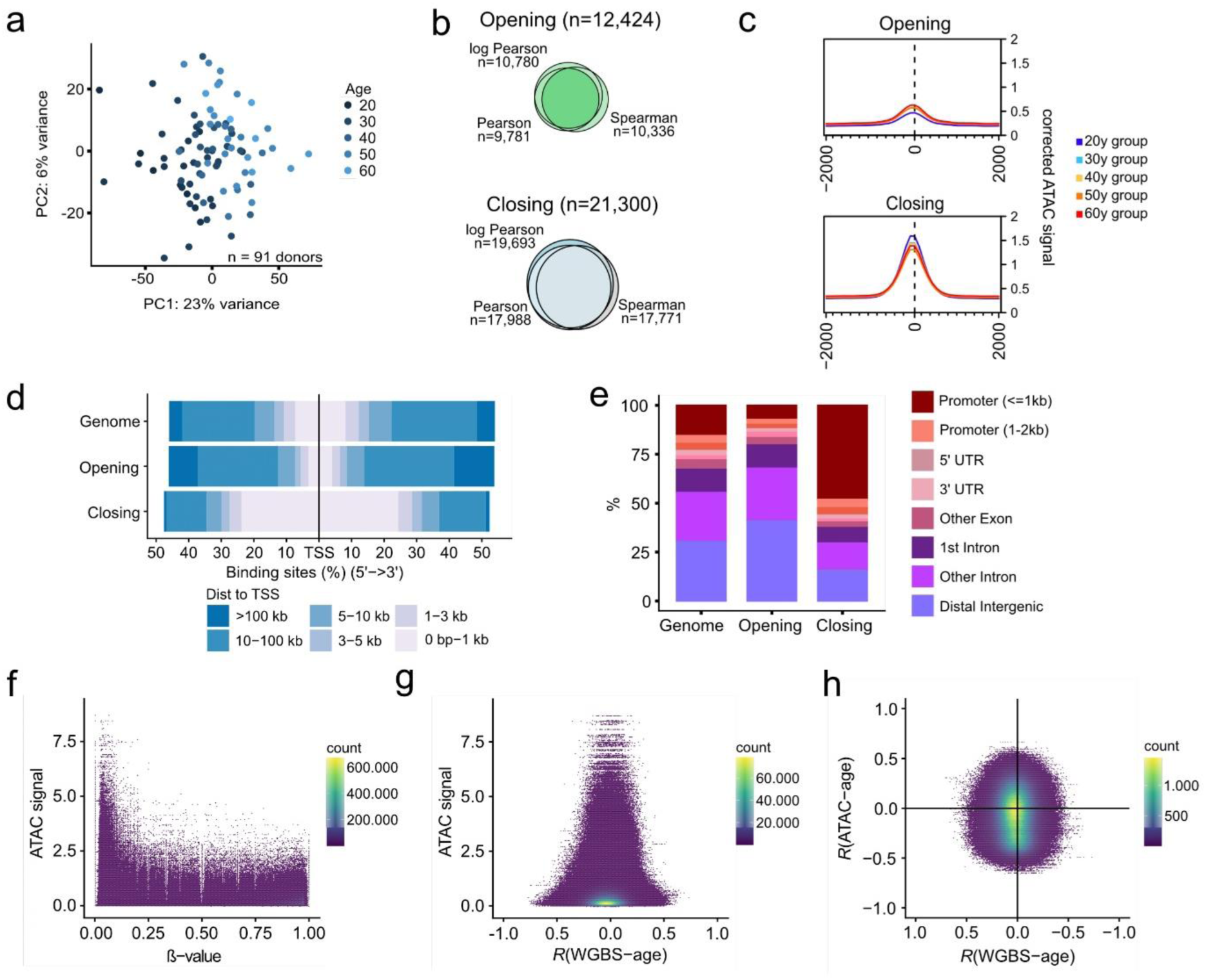
Age-associated chromatin accessibility dynamics identified by ATAC-seq. **a)** Principal component analysis (PCA) of 91 ATAC-seq samples based on the peak count matrix of age-associated open chromatin regions (OCRs), with samples colored according to age group. **b)** Age-associated closing and opening OCRs were identified using Pearson, log-transformed Pearson, or Spearman correlation with chronological age (|*R*| > 0.25; FDR-adjusted *P* < 0.05). The union of the three approaches comprised 21,300 closing and 12,424 opening OCRs. **c)** Aggregate chromatin accessibility profiles across opening and closing OCRs in different age groups (age categories in 20s, 30s, 40s, 50s, and 60s), shown within a ±2-kb window centered on each peak. **d,e)** Genomic distribution of all ATAC-seq peaks, opening OCRs, and closing OCRs relative to **d)** transcription start sites (TSSs) and **e)** gene features. **f)** Chromatin accessibility, quantified from ATAC-seq signal in fixed 500-bp genomic windows, was compared with WGBS-derived DNAm levels at the corresponding CpG sites to assess the relationship between chromatin accessibility and DNA methylation (representative example from a 30-year-old donor). **g)** Chromatin accessibility in the same representative donor was compared with the age-associated DNAm changes at individual CpG sites, quantified as the Spearman correlation coefficient across all WGBS donors. Relationship between age-associated changes in chromatin accessibility (Spearman correlation coefficient within ATAC-seq peaks) and age-associated DNA methylation changes at the corresponding CpG sites identified by WGBS.

We next investigated the relationship between age-associated chromatin accessibility changes and DNAm alterations. As expected, ATAC-seq signal intensity was substantially higher at unmethylated CpGs than at methylated CpGs (Fig. 2f). However, CpGs with higher age-associated correlation coefficients did not occur preferentially at regions with high or low ATAC-seq signal (Fig. 2g). Furthermore, when restricting the analysis to CpGs located within OCRs, we found no consistent relationship between age-associated changes in DNAm and changes in chromatin accessibility (Fig. 2h).

While we did not find consistent changes in chromatin accessibility at age-associated CpGs, the age-hypomethylated CpGs were rather in regions of lower baseline accessibility than hypermethylated CpGs (Supplementary Fig. 3d). Only a small proportion of age-associated DNAm alterations overlapped with age-associated OCRs, including approximately 3% of significant age-associated CpGs and 10% of DMRs. Although limited in number, the overlapping regions included several biologically relevant aging-associated loci, including *ELOVL2*, *GATA2*, *FOXP1,* and the senescence-associated *IGFBP7/IGFBP7-AS1* locus (Supplementary Fig. 3e). Thus, while individual genomic regions show coordinated age-associated changes in both DNAm and chromatin accessibility, these events appear to represent exceptions rather than a general feature of epigenetic aging.

### Association with age-associated remodeling of histone modifications

To determine whether age-associated changes in DNAm or chromatin accessibility are linked to specific chromatin features, we profiled CTCF occupancy and key histone modifications by ChIP-seq in PBMCs from five young donors (21–26 years) and five older donors (66–78 years). The analyzed histone marks included the active promoter mark H3K4me3, the active enhancer mark H3K27ac, the enhancer-associated mark H3K4me1, and the repressive modifications H3K9me3 and H3K27me3. Differential peak analysis identified age-associated gains and losses for all analyzed chromatin features (Supplementary Fig. 4).

We next examined whether age-associated DNAm changes occurred within genomic regions marked by altered CTCF occupancy or histone modifications. The 33 age-hypermethylated CpGs showed enrichment for CTCF and histone modification peaks, likely reflecting their preferential localization within promoter-associated regions. In contrast, the 585 age-hypomethylated CpGs showed no enrichment for any of the chromatin features. Importantly, ChIP-seq signal intensity at these regions did not differ between young and old donors, indicating that individual age-associated CpGs generally do not coincide with age-dependent changes in CTCF binding or histone modification (Fig. 3a).

**Figure 3.**
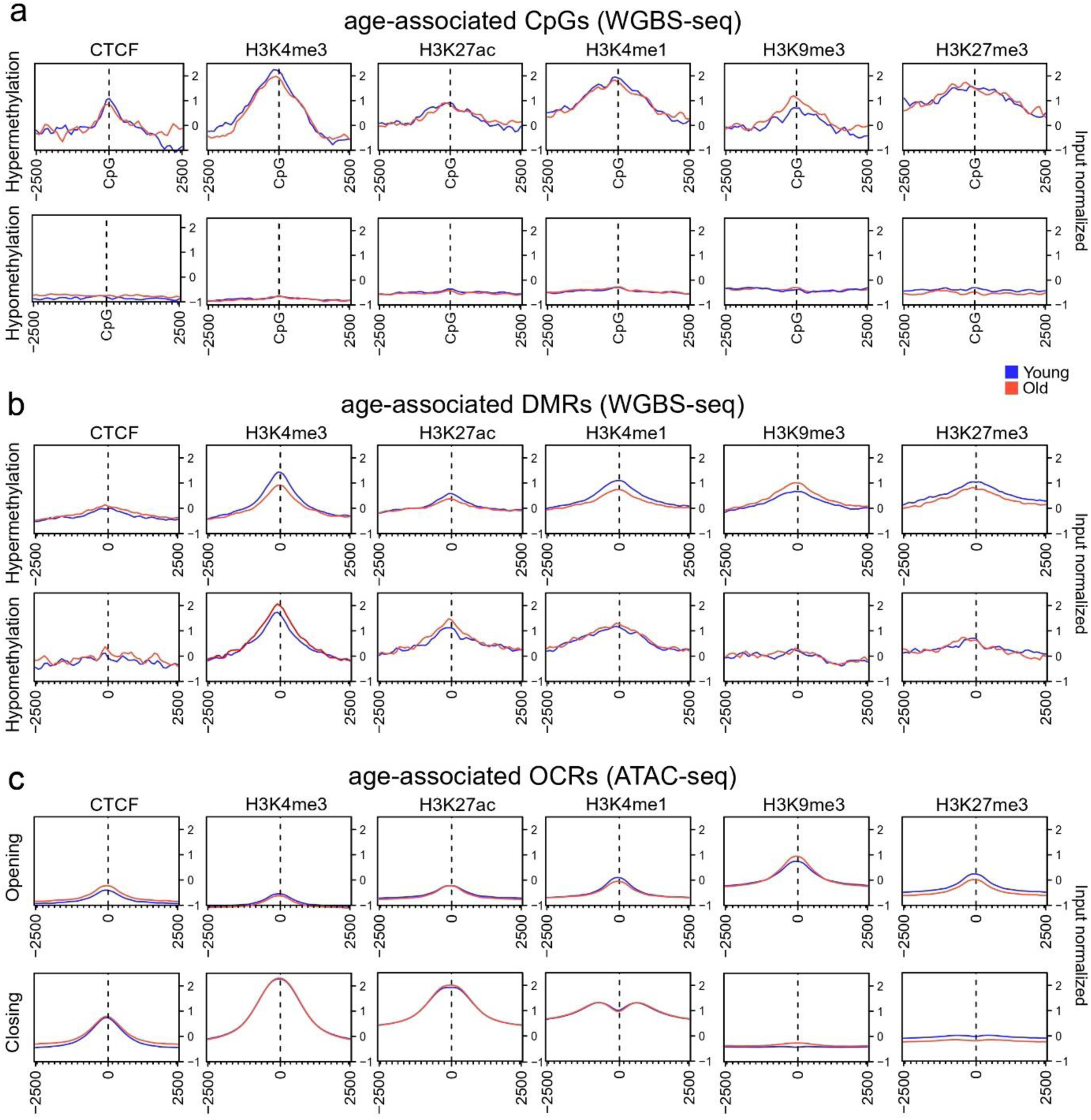
CTCF occupancy and histone modifications and at age-associated genomic regions. **a)** Aggregate ChIP-seq signal profiles for CTCF, H3K4me3, H3K27ac, H3K4me1, H3K9me3, and H3K27me3 in five young (21–26 years) and five older (66–78 years) donors, plotted within a ±2.5-kb window centered on the 585 age-hypomethylated and 33 age-hypermethylated CpG sites. **b)** Corresponding aggregate ChIP-seq signal profiles centered on the 268 age-hypermethylated and 31 age-hypomethylated differentially methylated regions (DMRs). Aggregate ChIP-seq signal profiles centered on the peaks of age-associated opening and closing open chromatin regions (OCRs) identified by ATAC-seq.

In contrast, hypomethylated DMRs showed enrichment for specific histone modifications. Moreover, hypo- and hypermethylated DMRs displayed opposing relationships at active chromatin marks H3K4me3 and H3K27ac, with in tendency, loss of activation marks at hyper-DMRs and *vice versa* (Fig. 3b). These findings suggest that regional methylation changes occur preferentially within specific chromatin contexts, whereas this is hardly observed for individual age-associated CpGs.

We next investigated whether age-associated chromatin accessibility changes were linked to altered histone landscapes or CTCF occupancy. Opening OCRs were enriched at CTCF binding sites and all histone modifications, particularly at the repressive marks H3K9me3 and H3K27me3. In contrast, closing OCRs show association with CTCF binding sites, as well as pronounced association with active chromatin marks K3K4me3, H3K27acm and H3K4me1, but not with H3K9me3 and H3K27me3. However, similar to the DNAm analysis, age-associated changes in OCRs were hardly accompanied by corresponding changes in CTCF occupancy or histone modification levels between young and old donors (Fig. 3c).

### Integration of age-associated epigenetic changes with chromatin states

To further investigate the relationship between age-associated epigenetic alterations and the broader chromatin landscape, we integrated the ChIP-seq datasets using a multivariate hidden Markov model (ChromHMM), which enables genome-wide annotation of combinatorial chromatin states based on the spatial distribution of histone modifications and CTCF occupancy ^31^. Using a 12-state model, we generated chromatin-state maps for young and older donors that showed moderate differences (Supplementary Fig. 5; Supplementary Table 5).

The 585 age-hypo CpGs were predominantly located within quiescent chromatin states, whereas the 33 age-hypermethylated CpGs showed enrichment for regulatory chromatin states, including bivalent/poised promoters, primed enhancers, and Polycomb-associated poised regions. This observation is consistent with the enrichment of age-associated hypermethylation with binding of polycomb repressive complex 2 (PRC2) ^32^. Despite these differences in baseline chromatin context, the chromatin-state annotations did not differ between young and old samples at the age-associated CpGs, indicating that at least the most prominent DNAm changes observed in our WGBS dataset were not driven by changes in chromatin state during aging (Fig. 4a,b).

**Figure 4.**
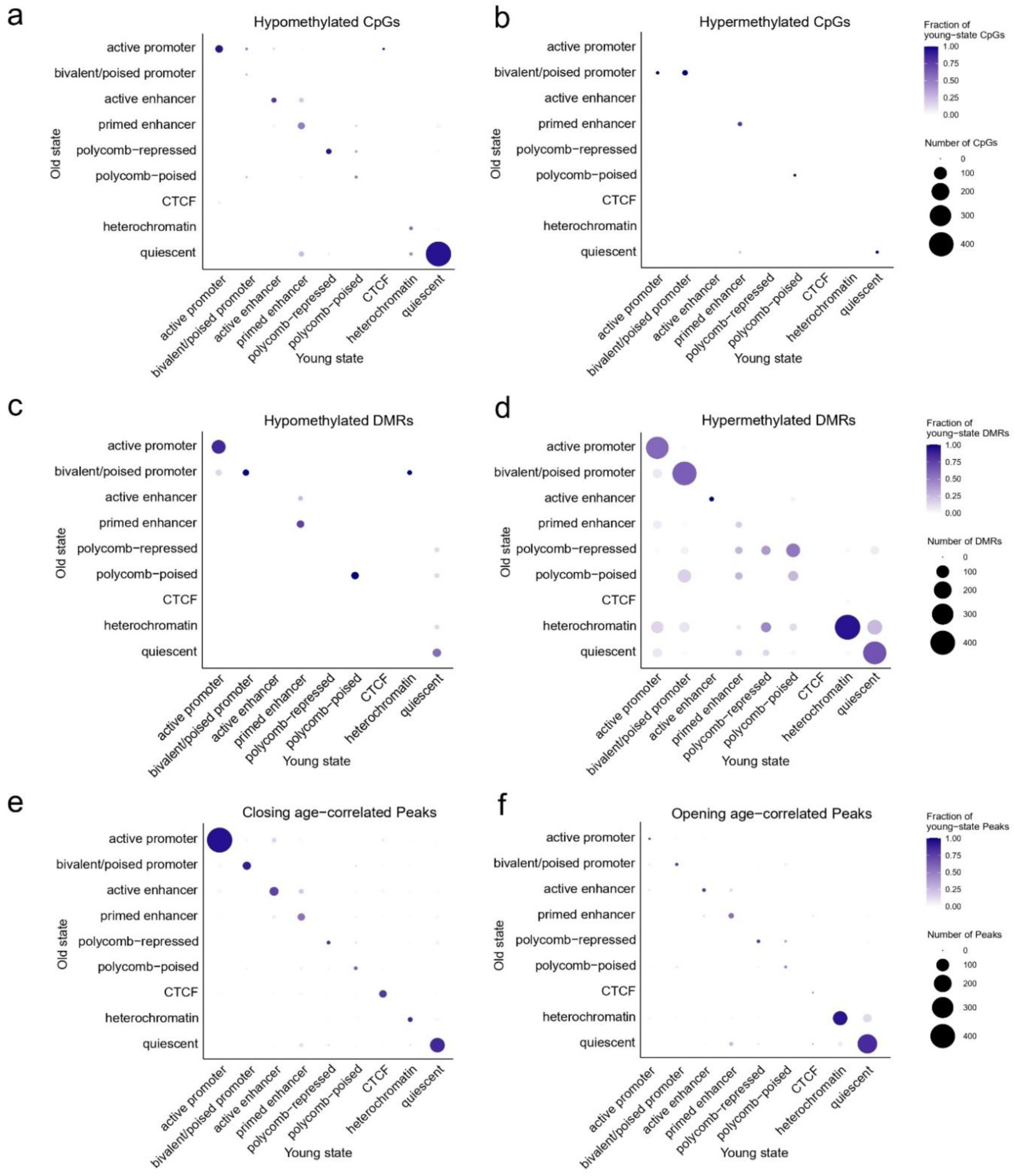
Chromatin states at age-associated genomic regions. **a,b)** Chromatin states were annotated using ChromHMM ^31^ and evaluated at age-associated CpG sites. The 585 age-hypomethylated CpGs were preferentially located within quiescent and heterochromatic states (**a**), whereas the 33 age-hypermethylated CpGs were enriched in promoter-associated chromatin states (**b**). Overall, the chromatin state assignments at these loci remained largely unchanged between young (21–26 years) and older (66–78 years) donors. **c,d)** Chromatin states at age-associated DMRs. The 31 age-hypomethylated DMRs were enriched in active promoter-associated chromatin states and showed little change in chromatin state with age (**c**). In contrast, the 268 age-hypermethylated DMRs were enriched in active promotor as well as quiescent and heterochromatic states in young donors and exhibited a gradual shift toward more regulatory chromatin states with aging (**d**). **e,f)** Chromatin states at age-associated changes in chromatin accessibility. The 21,300 closing OCRs were enriched in active and bivalent promoter-associated states (**e**), whereas the 12,424 opening OCRs were preferentially associated with quiescent and heterochromatic states (**f**). Overall, chromatin state assignments remained largely stable despite age-associated changes in chromatin accessibility.

Age-hypomethylated DMRs were enriched in active promoter states, whereas hypermethylated DMRs were associated with multiple chromatin environments, including active promoters, bivalent/poised promoters, heterochromatin, and quiescent chromatin states. Notably, in regions with hypermethylated DMRs, some changes in chromatin-state annotation occurred between young and older donors: regions classified predominantly as active promoters in young donors shifted toward heterochromatic and quiescent states in older donors. These findings suggest that, unlike individual age-associated CpGs, a subset of regional methylation changes occurs in the context of broader chromatin-state remodeling during aging (Fig. 4c,d).

Finally, we examined whether age-associated alterations in chromatin accessibility were accompanied by changes in chromatin states. Closing OCRs (21,300 regions) were enriched in active promoter states, whereas opening OCRs (12,424 regions) showed greater representation within heterochromatic and quiescent chromatin states. However, the chromatin-state composition of these regions was largely unchanged between young and older donors (Fig. 4e,f).

## Discussion

Aging is clearly associated with specific changes in methylome, chromatin accessibility, and histone code. However, our integrative analysis revealed only moderate association between the different epigenetic layers, particularly between age-associated DMRs and chromatin state changes but this was much less pronounced than would be anticipated if age-associated DNAm changes were primarily caused by chromatin accessibility or histone modifications. Previous studies have suggested links between age-associated DNAm changes and chromatin states ^33–35^, and age-related alterations in chromatin accessibility and histone modifications have been associated with transcriptional changes during aging ^7^. However, these studies generally focused on individual epigenetic layers or examined chromatin states without directly comparing age-dependent changes between young and old individuals.

A major resource generated by this study is the high-quality WGBS dataset from 120 individuals spanning a broad age range. Despite the widely described phenomenon of age-associated global hypomethylation ^16^, our genome-wide analysis revealed only a very modest reduction in mean DNAm levels with age, without preferential enrichment at endogenous retroelements or other repetitive sequences. This might be due to differences in cell type composition or malignant transformation in other studies ^15^.

The finding that the age-associated changes in different epigenetic layers seem to occur largely independent may support the notion that these processes rather resemble epigenetic drift than a fully coordinated remodeling program ^36^. Mathematical models have suggested that stochastic processes may account for substantial features of DNAm-based aging clocks ^37,38^. Whether similar stochastic mechanisms contribute to age-associated changes in chromatin accessibility and histone modifications remains an important open question. Either way, the concept of epigenetic drift does not exclude regulated components of aging ^39^.

Several limitations should be considered. We focused on blood samples to facilitate comparison with existing epigenetic clock studies and publicly available datasets. However, age-associated changes in leukocyte composition can influence epigenetic profiles ^40^. Another limitation is that DMR analyses and ChIP-seq profiling were performed in subsets of donors, limiting statistical power for detecting subtle continuous age-dependent changes. Last but not least, it is conceivable that the interplay of the different epigenetic aging trajectories would be more apparent within individual cells and therefore multi-omic single-cell sequencing analysis should further address this question in the future ^36^.

## Conclusion

Our integrative genome-wide analysis demonstrates that aging is associated with widespread alterations across multiple epigenetic layers, including methylome, chromatin accessibility, and histone modifications. Although these changes occur within characteristic chromatin contexts, they show surprisingly limited overlap and association. Rather than representing a single hierarchical aging program, different components of the epigenome appear to undergo partially overlapping but apparently largely independent age-associated changes.

## Methods

### Blood samples

Peripheral blood samples were obtained from 120 healthy adult donors (Supplemental Table 1) after written informed consent. All procedures were performed in accordance with the Declaration of Helsinki and approved by the ethics committees of RWTH Aachen University (EK 206/09 and EK 041/15). Whole blood was collected into EDTA-coated Vacutainer tubes and stored at 4°C until processing within several hours of collection. For ATAC-seq and ChIP-seq analysis PBMCs were isolated by density gradient centrifugation using Pancoll (Pan-Biotech).

### Methylome analysis

Genomic DNA was isolated from blood using the Blood & Tissue Kit (Qiagen) and purity was assessed using a NanoDrop spectrophotometer (Thermo Fisher Scientific). A total of 100 ng of genomic DNA was acoustically sheared to a mean fragment size of ∼300 bp using a Covaris S2 sonicator and microTUBE AFA Fiber Pre-Slit Snap-Cap tubes (Covaris, #520045). Fragmented DNA was purified and concentrated using the DNA Clean & Concentrator-5 kit (Zymo Research, #D4013) and eluted in low-TE buffer. Bisulfite conversion of purified DNA was done with the EZ DNA Methylation-Gold kit (Zymo Research, #D5005) according to the manufacturer’s protocol. Libraries were prepared with the xGen Methyl-Seq DNA Library Prep kit (Integrated DNA Technologies, #10009824), using seven PCR amplification cycles and two final rounds of purification with AMPure XP beads (Beckman Coulter, #A63881). WGBS was performed at the Cologne Center for Genomics within the West German Genome Center (WGGC). Libraries were sequenced on an Illumina NovaSeq 6000 platform in paired-end 2 × 150 bp mode, generating approximately 440-1100 million read pairs per sample.

Adapter trimming and quality trimming were performed with cutadapt (v4.6; parameters: - quality-cutoff 20 –overlap 5 –minimum-length 25; Illumina TruSeq adapter clipped from both reads), followed by trimming of 10 and 5 nucleotides from the 5’ and 3’ end of the first read and 15 and 5 nucleotides from the 5’ and 3’ end of the second read. Trimmed reads were aligned to the human reference genome hg38 using BSMAP (v2.90; parameters: -v 0.1 -s 16 -q 20 -w 100 -x 1000 -S 1 -u -R) ^41^. A sorted BAM file was obtained and indexed using samtools (vX.X) with the ‘sort’ and ‘index’ commands. Duplicates were removed using GATK4 (v4.5.0.0) ^42^ MarkDuplicates with lenient validation, duplicate removal enabled, and coordinate-sorted input. Finally, methylation rates were called using mcall from the MOABS package (v1.3.9.6; default parameters) ^43^. All methylation analyses were restricted to autosomes, and only CpGs covered by at least 10 and at most 150 reads were considered for downstream analyses. Global methylation levels were derived using average (arithmetic mean) genome-wide methylation levels. Region-specific methylation was assessed by calculating average (arithmetic mean) methylation levels across features using bigWigAverageOverBed from UCSC tools for each sample, where a feature was only considered if at least three CpGs were covered within a region.

### Identification of age-correlated CpGs

Age-associated CpG sites were identified using Pearson correlation, Pearson correlation on log-transformed age, and Spearman correlation between methylation rates and chronological age. CpGs with fewer than 60 non-missing values across the 120 samples (i.e., <50% coverage) were excluded. P-values were adjusted for multiple testing using Benjamini-Hochberg, and CpGs were considered age-associated per method if the absolute correlation coefficient was > 0.6 and the adjusted p-value was < 0.05. The age-associated CpGs from Pearson, log-Pearson, and Spearman were combined to yield the union of age-correlated CpG sites.

### Differential methylation analysis

Differentially methylated regions (DMRs) were called using metilene (v0.2-8) ^44^ between 5 young and 5 old samples (indicated in Supplemental Table 1). DMRs were defined by an absolute minimum difference in methylation of 0.1 with a maximum distance of 300nt between CpGs within a DMR and a minimum of 10 CpGs per DMR and Bonferroni correction for multiple testing. DMRs were filtered for a significant q-value (q < 0.05) and classified as ‘Hypo DMRs’ and ‘Hyper DMRs’ based on the direction of change.

### Genomic Feature Annotation

CpG islands were obtained from the UCSC table browser (https://genome-euro.ucsc.edu/cgi-bin/hgTables) ^45^. CpG shores were defined as 2 kb regions flanking CGIs upstream and downstream; shelves were defined as 2 kb regions flanking the shores. Genomic regions (DMRs, peaks) or loci (CpGs) were annotated with respect to genomic features (promoter, 5’/3’ UTR, first exon, other exon, intron, and distal intergenic; in case of multiple overlaps, the assignment was based on precedence as defined by the given order) and distance to the nearest transcription start site using the ChIPseeker ^46^ and TxDb.Hsapiens.UCSC.hg38.knownGene package (v3.18.0). Repeat annotations were obtained from the RepeatMasker track via UCSC table browser ^45^, removing duplicate entries at identical genomic coordinates. CpG density was calculated as the fraction of CpG dinucleotides within 250 bp windows using a sliding window approach (50 bp step size). Annotations of solo-WCGW CpGs were obtained from https://zwdzwd.github.io/pmd.

### Epigenetic clocks

We evaluated eight epigenetic age predictors originally developed for Illumina BeadChip methylation array data. To apply these clocks to our WGBS data, CpGs with a sequencing coverage <5 were imputed on a per-sample basis using the BoostMe package (v0.1.0), with each sample represented as a bsseq object (v1.40.0). Imputed methylation values were capped at 1. Illumina probe IDs were mapped to genomic coordinates using the EPIC array manifest. Epigenetic ages were then estimated using the DNAmAge function from the methylclock package (v1.10.0) ^47^. Prediction accuracy was assessed by calculating the median absolute error (MAE; years) between chronological and predicted age and the Pearson correlation coefficient across all samples.

### Analysis of age-associated gene expression

To investigate the relationship between age-associated DNAm changes and gene expression, we analyzed publicly available PBMC RNA-seq data from 157 healthy donors aged 20–74 years (GSE193142) ^3^. Gene-level counts were normalized using the edgeR package (v4.4.2) with the trimmed mean of M-values (TMM) method. Following the quality control procedures described in the original study, 15 samples identified as outliers were excluded prior to downstream analyses ^3^. Age-associated gene expression was assessed by calculating Spearman correlations between normalized gene expression levels (counts per million, CPM) and donor age.

### ATAC-seq library preparation and preprocessing

ATAC-seq libraries were generated from 96 PBMC samples (donor age: 20–70 years) from individuals included in the DNA methylation cohort (Supplementary Table 1) using an adapted Omni-ATAC protocol ^48^. Briefly, approximately 60,000 freshly isolated PBMCs per donor were subjected to cell lysis, nuclear isolation, and transposition. Transposed DNA was purified and shipped on dry ice to the Cologne Center for Genomics within the West German Genome Center (WGGC) for library preparation and sequencing. Libraries were sequenced on an Illumina NovaSeq 6000 platform in paired-end 2 × 150 bp mode, generating approximately 60 million read pairs per sample.

Raw sequencing data were processed using the nf-core/atacseq pipeline (v2.0). Adapter trimming and quality filtering were performed with Trim Galore (v0.6.7), and high-quality reads were aligned to the GRCh38 human reference genome using BWA (v0.7.17). PCR duplicates were removed prior to peak calling. Sample quality was assessed using standard ATAC-seq metrics, including the fraction of reads in peaks (FRiP) ^49^, transcription start site (TSS) enrichment ^50^ calculated with ATACseqQC (v1.33), total mapped reads, peak number, and nucleosome periodicity. Five samples with extremely low FRiP scores (<0.01) were excluded from further analyses.

Accessible chromatin regions were identified using MACS2 (v2.2.7.1) with narrow peak calling and without read shifting or extension. Only autosomal peaks were retained for downstream analyses. To generate a consensus peak set, peaks from 18 high-quality samples (FRiP > 0.1) were merged using BEDTools (v2.30.0), retaining regions detected in at least two samples. Read counts were quantified across the consensus peak set using featureCounts from the Subread package (v2.0.1), producing a peak-by-sample count matrix. Peaks with fewer than 10 reads in at least 10 samples were removed, resulting in a final set of 239,599 consensus open chromatin regions (OCRs).

### ATAC-seq analysis

Bulk ATAC-seq count data were normalized using the trimmed mean of M-values (TMM) method implemented in edgeR (v4.4.2). Normalized counts were transformed to log2 counts per million (logCPM) using the voom function in limma (v3.54.2). Technical batch effects, sex, and differences in cell-type composition were regressed out using linear modeling in limma.

To estimate cell-type composition, we generated a chromatin accessibility signature matrix from our previously published single-cell multiomics dataset ^51^. Briefly, accessibility counts for cell-type-specific marker peaks were aggregated across all cells of each cell type (n = 10 donors) to generate pseudobulk profiles representing cell-type-specific chromatin accessibility. Cell-type proportions in the bulk ATAC-seq samples were then estimated using CIBERSORT (v0.1.0) ^52^ and DeconPeaker ^53^. To determine the optimal signature matrix, we evaluated different numbers of marker peaks per cell type using sample-specific pseudobulk profiles as validation data and compared deconvolution performance based on the mean squared error (MSE). Based on this benchmarking, a signature matrix comprising the top 500 marker peaks per cell type was used for all downstream analyses.

Age-associated changes in chromatin accessibility were assessed by calculating Spearman’s rank correlation, Pearson correlation, and Pearson correlation using log-transformed age between logCPM values and donor age. OCRs were considered age-associated if they exhibited an absolute correlation coefficient of at least 0.25 and an FDR-adjusted *P* value ≤ 0.05. OCRs were annotated to the nearest gene using ChIPseeker (v1.42.1) ^46^, with promoter regions defined as ±3 kb around the transcription start site (TSS).

To characterize genome-wide changes in chromatin accessibility independently of peak calling, we employed a fixed-bin approach. The genome was partitioned into non-overlapping 500 bp bins, excluding ENCODE blacklist regions ^54^, and only autosomal chromosomes were retained for downstream analyses. Reads overlapping each genomic bin were quantified using featureCounts (v2.20.0), generating a genome-wide count matrix. Raw counts were normalized using the trimmed mean of M-values (TMM) method implemented in edgeR (v4.4.2) and converted to counts per million (CPM). Technical batch effects were corrected using the ComBat_seq function from the sva package (v3.54.0). The resulting normalized accessibility profiles were integrated with the WGBS data for subsequent joint analyses.

### ChIP-seq analysis

Chromatin immunoprecipitation followed by sequencing (ChIP-seq) was performed on PBMCs from five young (21–26 years) and five older donors (66–78 years; Supplementary Table 1) to profile the histone modifications H3K4me1, H3K4me3, H3K9me3, H3K27ac, and H3K27me3, as well as CTCF binding. ChIP was performed using the ChIP-IT PBMC Kit (Active Motif, Carlsbad, CA) according to the manufacturer’s instructions. Briefly, PBMCs were crosslinked, lysed, and sonicated using a Covaris M220 focused ultrasonicator to generate chromatin fragments of 200–1,000 bp. For each immunoprecipitation, 30 μg of chromatin was incubated with antibodies against H3K27ac (ab4729, Abcam), H3K27me3 (ab6002, Abcam), H3K4me1 (ab8895, Abcam), H3K4me3 (ab8580, Abcam), H3K9me3 (ab8898, Abcam), or CTCF (61311, Active Motif). Following immunoprecipitation, crosslinks were reversed and DNA was purified.

Sequencing libraries were prepared and sequenced at the Cologne Center for Genomics within WGGC on an Illumina NovaSeq 6000 platform in paired-end 2 × 150 bp mode, generating approximately 20 million read pairs per sample. Raw sequencing data were processed using the nf-core/chipseq pipeline (v2.0.0). Adapter trimming and quality filtering were performed with Trim Galore, and reads were aligned to the GRCh38 human reference genome using BWA. PCR duplicates were removed prior to peak calling.

Peaks were identified using MACS2 (v2.2.7.1), applying narrow peak calling for CTCF, H3K4me3, and H3K27ac, and broad peak calling for H3K4me1, H3K9me3, and H3K27me3. For each chromatin feature, a consensus peak set was generated by merging overlapping peaks across samples using BEDTools. Differentially enriched regions between young and older donors were identified using THOR from the RGT suite (v1.0.0) ^55^, and *P* values were adjusted for multiple testing using the Benjamini–Hochberg procedure. Regions were considered significantly gained or lost if they exhibited an adjusted *P* value < 10^-20^ for histone modifications or < 0.01 for CTCF, together with an absolute log2 fold change > 0.5. Differential regions were annotated to the nearest gene using ChIPseeker (v1.42.1), with promoter regions defined as ±3 kb around the TSS.

### ChromHMM analysis

Chromatin states were inferred using ChromHMM (v1.27) ^31^ based on ChIP-seq data for five histone modifications (H3K4me1, H3K4me3, H3K9me3, H3K27ac, and H3K27me3), and CTCF. Aligned BAM files were binarized using the BinarizeBam module, with matched input controls used to account for background signal. Separate chromatin state models were learned for young and older donors using the LearnModel function with 12 states and a 200 bp bin size, generating genome-wide chromatin state annotations for each age group. Individual chromatin state segmentations were then generated using the MakeSegmentation function. To facilitate biological interpretation, the 12 chromatin states were assigned functional annotations based on their emission and transition probabilities together with enrichment for genomic features and previously described chromatin state signatures. Chromatin state annotations from young and older donors were subsequently intersected with age-associated CpGs, DMRs, and age-associated changes in chromatin accessibility to assess their genomic distribution.

### Visualizations

Principal component analyses (PCA) were calculated using the prcomp R function and visualized using ggplot2 (v3.5.2). Violin plots, volcano plots, bar plots, boxplots, scatter plots, line plots, 2D kernel density plots of methylation rate for individual CpG sites, and annotation summaries were generated using the ggplot2 package. Violin plots show the kernel density estimation with embedded boxplots indicating the median, interquartile range, and whiskers extending to 1.5× the interquartile range. Boxplots show the median (center line), interquartile range (box), and whiskers extending to 1.5× the interquartile range; individual observations are overlaid as points. Average bigWig tracks for WGBS and ATAC signal were generated using deepTools bigwigAverage (v3.5.5) prior to visualization. Browser shots were generated using the Integrated Genomics Viewer (IGV), and show averaged methylation rates across 5 young and 5 old samples (indicated in Supplemental Table 1) and averaged ATAC profiles across the 5 youngest and 5 oldest ATAC samples. Heatmaps and average profile tracks were created using the EnrichedHeatmap package (v1.34.0) by applying the normalizeToMatrix function, centered on the respective genomic region and extensions up- and downstream as shown. For ATAC and ChIP, normalizeToMatrix was applied to each sample’s bigWig track separately and resulting matrices were averaged within each group. For WGBS, precomputed average methylation tracks per age group were used, generated as described above. Venn diagrams were generated using the eulerr package (v7.0.4). Ridge plots were generated using the ggridges package (v0.5.4), with the median indicated by a vertical line.

## Additional Information

## Supporting information

Suppemental Table 5

## Acknowledgments

We thank the Cologne Center for Genomics within the West German Genome Center (WGGC) for library preparation and sequencing of the datasets generated in this study. This research was specifically supported by the Deutsche Forschungsgemeinschaft (GE 2811/5-1, to I.C.; ME 3449/4-1 to A.M.; and WA 1706/14-1 and SFB 1506-1/450627322 to W.W.), and also by the Else Kröner-Fresenius Stiftung (2025-EKSE.35 to W.W.), and the José Carreras Foundation (DJCLS 03 R/2024 to W.W.).

## Author contributions

M.S., R.K., and M.S. performed bioinformatic analysis. G.F., and D.P. carried out the experiments. H.K. contributed viable materials. A.M., I.C., H.K, and W.W. contributed to the experimental design and supervised the analysis. W.W. wrote the initial draft of the manuscript. All authors revised and approved the final version of the manuscript.

## Supplementary tables

**Supplementary table 1. Sample overview**. This Excel table summarizes all samples included in the study, including sample ID, age, sex, and whether each sample was included in WGBS, DMR analysis, ChIP-seq, ATAC-seq, or single-cell multi-omics analyses. For WGBS samples, the table reports the total number of sequencing reads, mapped reads, the number of CpGs with sufficient coverage, and the mean DNA methylation level. For ATAC-seq samples, the fraction of reads in peaks (FRiP) score and transcription start site (TSS) enrichment score are provided.

**Supplementary table 2. Age-associated CpGs identified by WGBS.** This Excel table lists the 618 CpG sites that showed a significant association with age based on Pearson, log-transformed Pearson, or Spearman correlation analyses. The table includes chromosome, genomic position, Pearson correlation coefficient, adjusted Pearson *P* value, log-transformed agePearson correlation coefficient, adjusted log-transformed Pearson *P* value, Spearman correlation coefficient, and adjusted Spearman *P* value.

**Supplementary table 3. Age-associated DMRs identified by WGBS.** This Excel table summarizes the 299 significantly age-associated DMRs identified using Pearson, log-transformed Pearson, or Spearman correlation analyses. The table includes chromosome, start position, end position, region width, mean DNA methylation difference (old – young), and adjusted *P* value.

**Supplementary table 4. Age-associated chromatin accessibility changes.** This Excel table summarizes the 12,346 chromatin regions showing increased accessibility (“opening”) and the 21,300 regions showing decreased accessibility (“closing”) with age. Regions were identified using Pearson, log-transformed age Pearson, or Spearman correlation analyses. The table includes chromosome, start position, end position, Pearson correlation coefficient and FDR, Spearman correlation coefficient and FDR, and log-transformed Pearson correlation coefficient and FDR.

**Supplementary table 5. ChromHMM annotation of genomic regions.** This Excel table provides ChromHMM annotations for the genomic regions analyzed in this study. For each region, the table reports the corresponding ChromHMM chromatin state assignment in young and older donors based on the reference annotation, enabling functional interpretation of age-associated DNAm and chromatin accessibility changes.

## Supplementary Figures

**Supplementary Figure 1.**
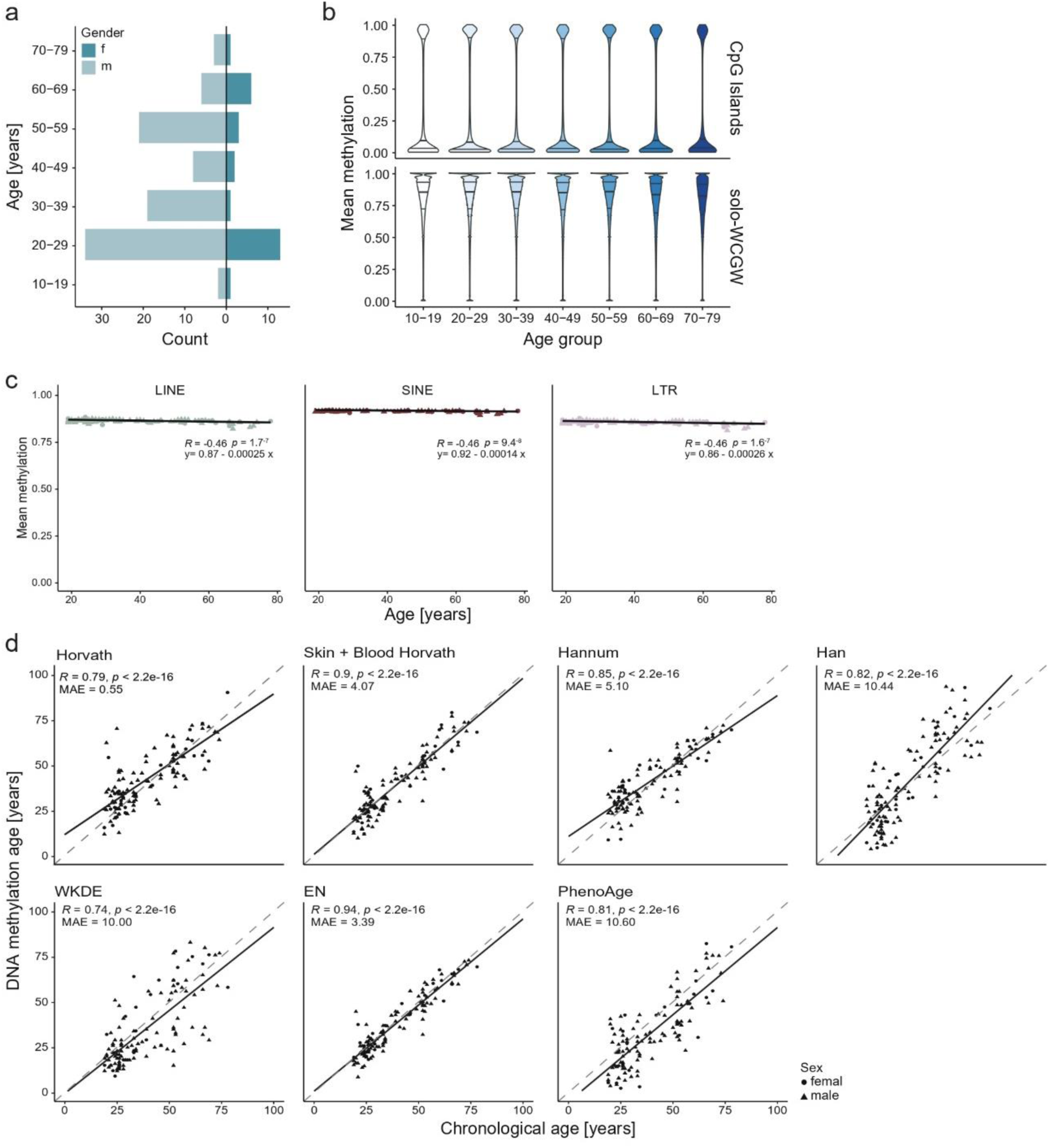
Age-associated DNA methylation loss and epigenetic age prediction. **a)** Age and sex distribution of the 120 blood donors included in the WGBS cohort. **b)** Age-associated loss of DNAm was slightly more pronounced at CpGs flanked by an A or T nucleotide (“W”) on both sides (solo-WCGW CpGs), which have previously been reported to be particularly susceptible to stochastic methylation loss during aging ^15^. **c)** Mean DNAm levels within repetitive elements declined modestly but significantly with age, including long interspersed nuclear elements (LINEs; β = –0.00025, *P* = 1.7 × 10⁻⁷), short interspersed nuclear elements (SINEs; β = –0.00014, *P* = 9.4 × 10⁻⁸), and long terminal repeat (LTR) retrotransposons (β = –0.00026, *P* = 1.6 × 10⁻⁷). Overall, repetitive elements exhibited a gradual age-associated hypomethylation comparable to the genome-wide average. **d)** Epigenetic age predictions were generated using the Horvath clock (353 CpGs) ^20^, Skin & Blood clock (391 CpGs) ^21^, Hannum clock (71 CpGs) ^22^, Han clock (65 CpGs) ^23^, PhenoAge (513 CpGs) ^23^, weighted kernel density estimation (WKDE; 27 CpGs) (^25^, and the elastic net (EN) clock (514 CpGs) ^26^. Pearson’s correlation coefficient (*R*) and the median absolute error (MAE) between predicted and chronological age are indicated

**Supplementary Figure 2.**
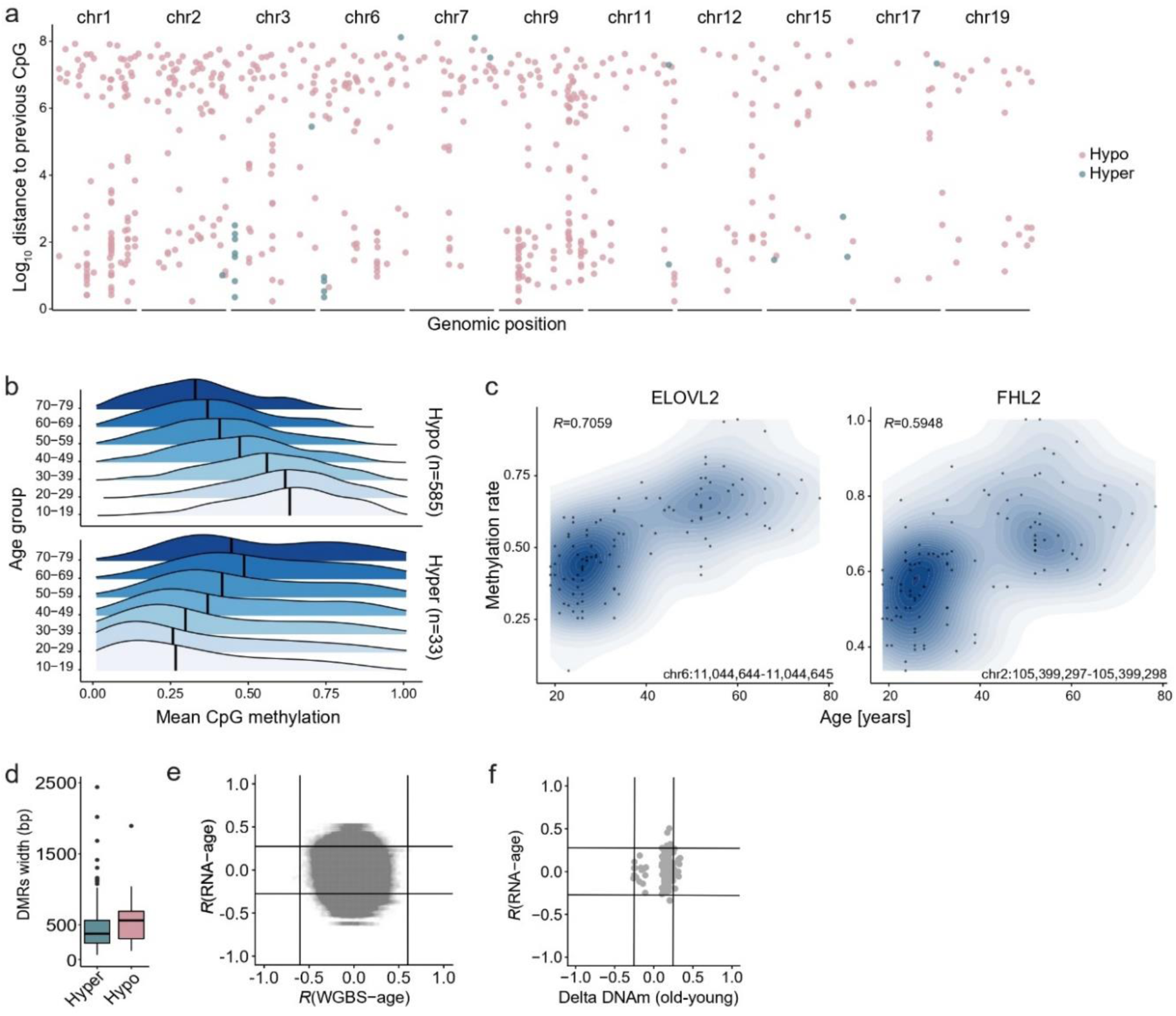
Genomic distribution and functional relevance of age-associated DNAm changes. **a)** Genome-wide distribution of the 585 age-hypomethylated and 33 age-hypermethylated CpG sites. Age-associated CpGs were broadly distributed across the genome, with several clusters of hypomethylated CpGs located within or near *ZBTB38, STAT4, MEIS1, SORBS3, TBX3, KLF5, CDH8*, and *ELOVL2*, whereas multiple hypermethylated CpGs were identified within *SEMA3F* and an intergenic region adjacent to *C6orf52*. **b)** Distribution of DNAm levels at the 33 age-hypermethylated (top) and 585 age-hypomethylated (bottom) CpG sites across different age groups. DNAm levels displayed substantial variability while shifting progressively toward higher or lower methylation with age, respectively. Although many of these CpGs exhibited intermediate methylation levels, they did not preferentially converge toward 50% methylation. **c)** Representative two-dimensional kernel density plots illustrating the relationship between DNAm and chronological age at the age-associated loci *ELOVL2* and *FHL2*, two well-established epigenetic aging markers ^23,28^ (Pearson correlation with age is indicated). **d)** Distribution of the widths of age-associated DMRs, including 268 age-hypermethylated and 31 age-hypomethylated DMRs. **e)** To assess whether age-associated promoter methylation was associated with transcriptional changes, public RNA-seq data (GSE193142) ^3^ were analyzed. Spearman correlation coefficients between normalized gene expression levels (counts per million, CPM) and donor age were compared with the corresponding age-associated CpGs. Overall, age-associated promoter methylation showed little association with age-related changes in gene expression. **f)** Similarly, age-associated gene expression changes were compared with methylation changes at DMRs, quantified as the difference in mean DNAm between the five young and five older donors. No consistent association was observed between differential methylation and corresponding gene expression changes.

**Supplementary Figure 3.**
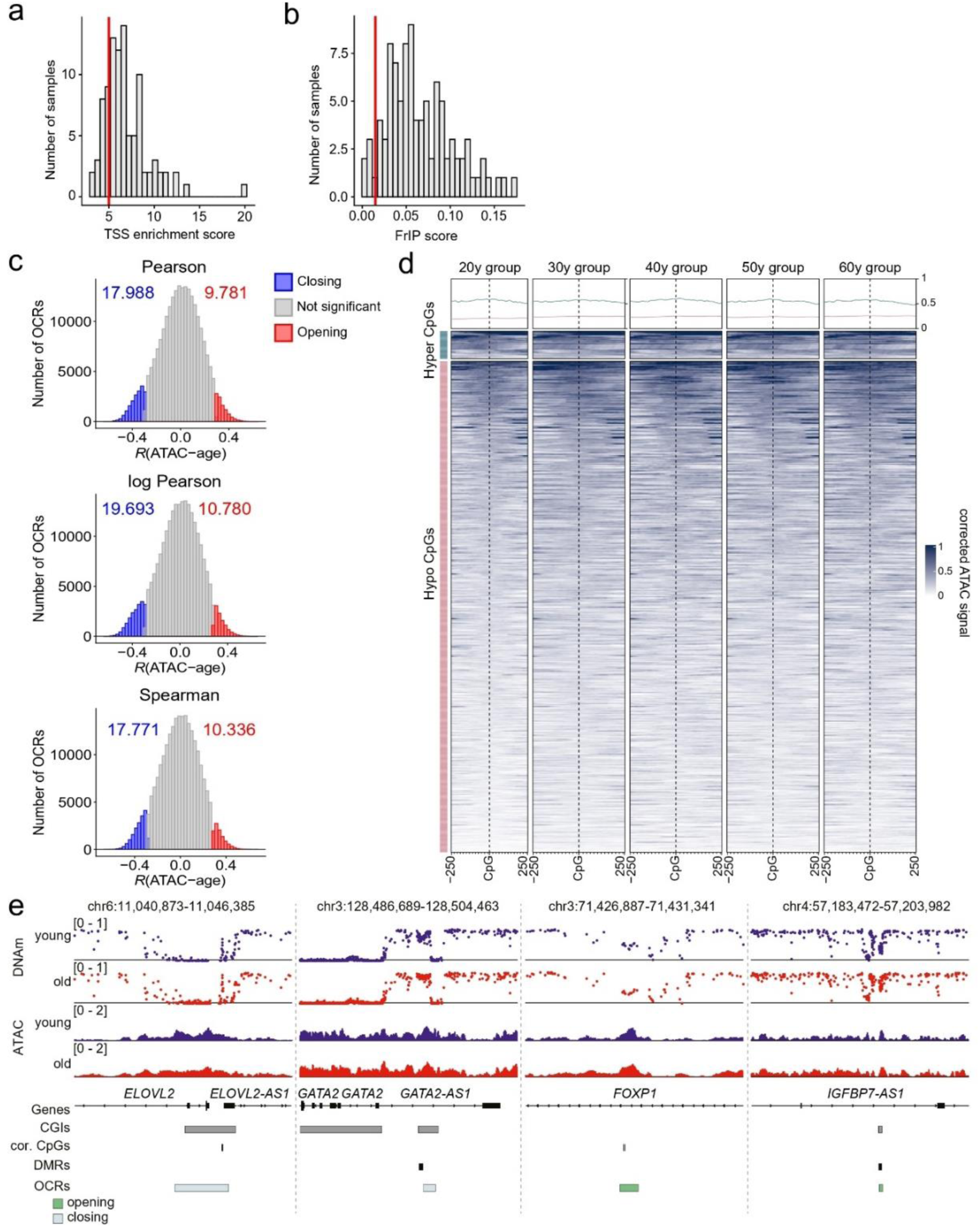
Age-associated changes in chromatin accessibility. **a,b)** Quality assessment of the 96 ATAC-seq libraries based on **a)** transcription start site (TSS) enrichment scores and **b)** the fraction of reads in peaks (FRiP) scores. Five samples were excluded from downstream analyses because of low FRiP scores. **c)** Distribution of correlation coefficients between ATAC-seq peak accessibility and chronological age calculated using Pearson, log-transformed Pearson, and Spearman correlation analyses. Age-associated opening and closing OCRs were defined by an FDR < 0.05 and correlation coefficients of *R* > 0.25 (red) and *R* < −0.25 (blue), respectively. **d)** Heatmaps of normalized ATAC-seq signal within a ±250-bp window centered on the union set of age-hypermethylated and age-hypomethylated CpG sites. Line plots above each heatmap show the corresponding average accessibility profiles. Samples are grouped into decade-based age categories (20s, 30s, 40s, 50s, and 60s). **e)** Exemplary IGV plots of the genomic regions in *ELOVL2*, *GATA2*, *FOXP1,* and *IGFBP7/IGFBP7-AS1* locus. DNAm values and ATAC-signals are depicted for five young and five old donors. The location of CpG islands (CGIs), age-associated CpGs (cor. CpGs), age-associated differentially methylated regions (DMRs), and opening/closing open chromatin regions (OCRs) are indicated.

**Supplementary Figure 4.**
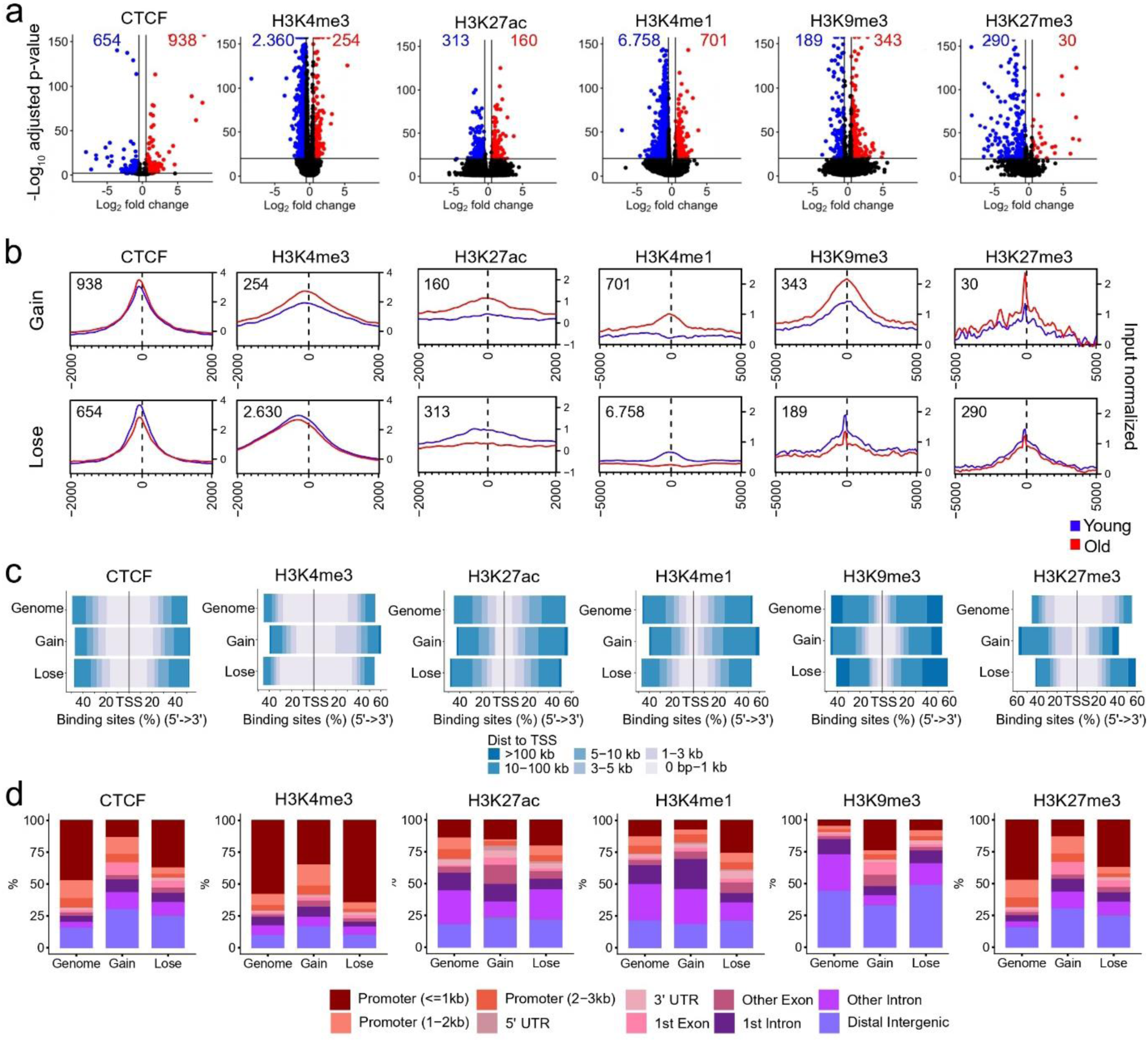
Age-associated changes in CTCF occupancy and histone modifications. **a)** Volcano plots showing differential CTCF occupancy and histone modification signals (H3K4me3, H3K27ac, H3K4me1, H3K9me3, and H3K27me3) between five young (21–26 years) and five older (66– 78 years) donors. Differential ChIP-seq peaks were identified using the THOR pipeline ^55^. Significant changes were defined as an adjusted *P* < 10^-20^ for histone modifications or an adjusted *P* < 0.01 for CTCF, together with an absolute log₂ fold change > 0.5. Regions with increased and decreased signal in older donors are shown in red and blue, respectively. **b)** Aggregate ChIP-seq signal profiles centered on regions exhibiting age-associated gains or losses in CTCF occupancy or histone modifications identified in (within ±2 or ±5 kb windows). The number of differential peaks is indicated for each chromatin feature. **c,d)** Genomic distribution of all ChIP-seq peaks and age-associated gained and lost peaks relative to **c)** transcription start sites (TSSs) and **d)** gene features.

**Supplementary Figure 5.**
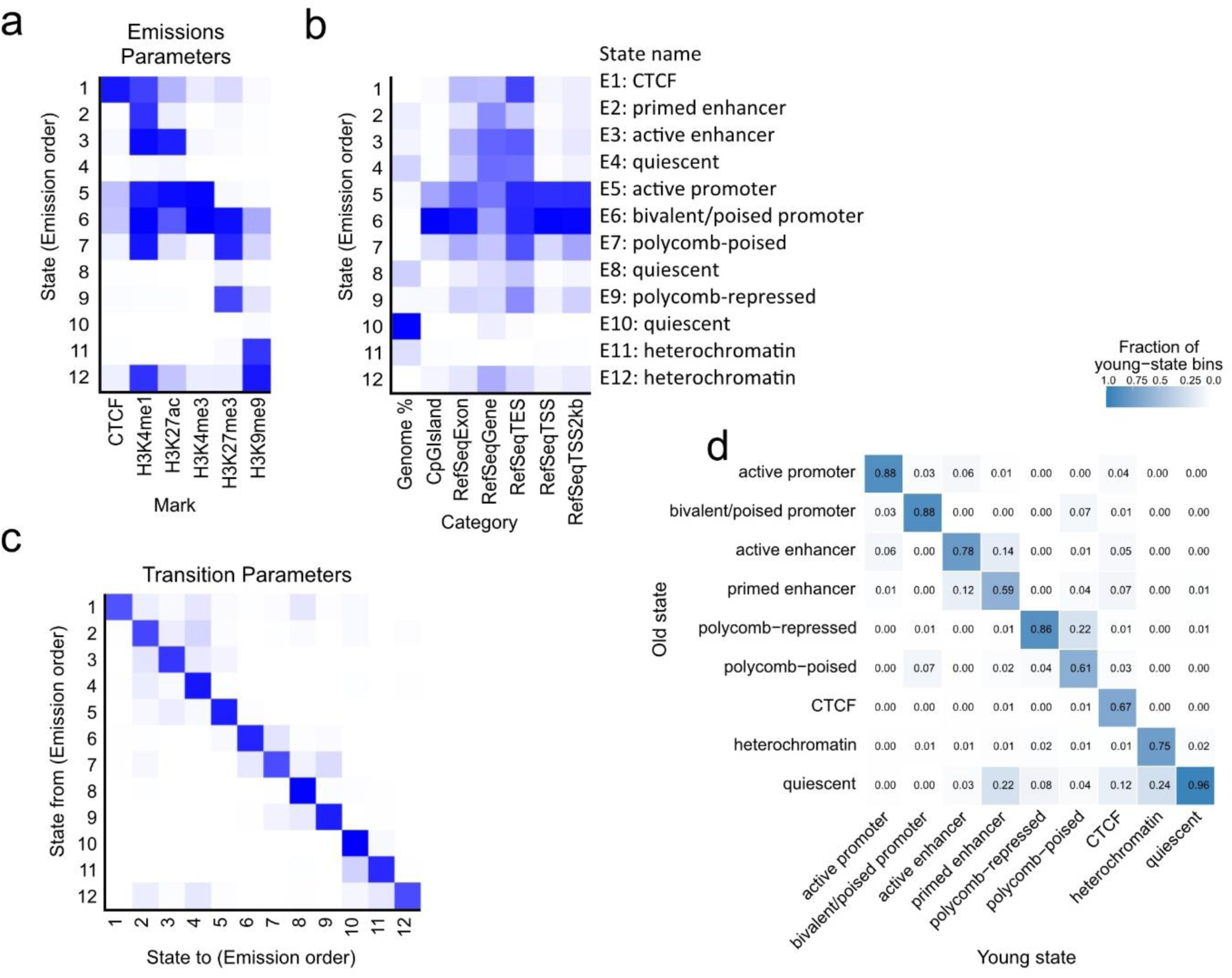
ChromHMM-based chromatin-state annotation in young and older donors. **a)** Emission probability heatmap of the 12-state ChromHMM model ^31^ trained on combined PBMC datasets from five young (21–26 years) and five older (66–78 years) donors. The model was generated using ChIP-seq data for CTCF, H3K4me3, H3K27ac, H3K4me1, H3K9me3, and H3K27me3. Darker blue shading indicates a higher emission probability of the respective chromatin feature within a given state. **b)** Functional annotation of the 12 ChromHMM states based on their enrichment for genomic features. Chromatin states were assigned to functional categories, including active promoter, enhancer, Polycomb-repressed, CTCF-associated, heterochromatin, and quiescent states. **c)** Transition probability heatmap of the 12-state ChromHMM model. Darker colors indicate higher transition probabilities between chromatin states. The pronounced diagonal enrichment reflects high self-transition probabilities, supporting the robustness and spatial continuity of the inferred chromatin-state annotations. **d)** Comparison of chromatin-state annotations between young and older PBMC samples. The heatmap shows the fraction of genomic bins shared between chromatin states obtained from age group-specific genome segmentations using the trained 12-state ChromHMM model. Strong diagonal enrichment indicates high concordance of chromatin-state assignments between age groups, whereas off-diagonal enrichment represents age-associated chromatin-state transitions.

